# Assessment of DNA methylation from a single genomic region of ELOV2 is sufficient to predict chronological age

**DOI:** 10.1101/2024.12.10.627662

**Authors:** Bowen Zhu, Dean Li, Guojing Han, Xue Yao, Hongqin Gu, Tao Liu, Linghua Liu, Jie Dai, Isabella Zhaotong Liu, Yanlin Liang, Jian Zheng, Zheming Sun, He Lin, Wenyuan Wang, Nan Liu, Haidong Yu, Meifang Shi, Gaofang Shen, Lefeng Qu

## Abstract

Estimation of chronological age is particularly informative in forensic contexts. Assessment of DNA methylation status allows for the prediction of age, though the accuracy and ease of manipulation may vary across different models. In this study, we started with a carefully designed discovery cohort recruiting more elderly subjects than other age categories, to diminish the effect of epigenetic drifting. We analyzed DNA methylation from a single genomic region of ELOV2, which was sufficient to construct an age-prediction model comprising 15 CpG sites. This model is further validated by an independent cohort as well as a multi-center test using trace dried bloodstains. The nature of our analytical pipeline, when combined the assessment of a single genomic locus with high-throughput sequencing, can easily be scaled up with low cost. Taken together, we propose a new age-prediction model featuring accuracy, ease of manipulation, high-throughput, and low cost. This model can be readily applied in both classic and newly emergent forensic contexts that require age estimation.

## Introduction

Forensic DNA phenotyping exploits not just genetic polymorphisms but also epigenetic modifications, i.e., DNA methylation, to draw a bio sketch of an unknown subject [1]. DNA methylation is a methyl group on the cytosine (C) followed by guanine (G) that is commonly referred to as CpG site, where the p stands for the phosphodiester bond between the two nucleotides. Recent studies have reported models that estimated chronological age from DNA methylation status, i.e., the ratio between methylated and non-methylated CpG forms. However, their prediction accuracy was hugely varied, usually deviating 3 to 5 or more years from the actual age [2–4]. Moreover, these models profiled DNA methylation from multiple genomic loci by low-throughput assays, i.e., pyrosequencing or PCR-based SNaPshot, thus limiting their ability to handle a large quantity of samples [5, 6].

Mounting evidence has associated aging and diseases with loss of fidelity in epigenetic drift of DNA methylation, i.e., stochastic changes of methylation status [7–9]. The currently available age-prediction models involved the use of five to eight CpG sites from distinct genomic regions and consequentially their actual performance could be perturbed by aging and pathophysiological conditions [4, 5, 10]. Indeed, a substantial decline of prediction accuracy in the elder population has been documented in previous models [5]. Notably, regardless of human ethical groups and applicable body fluids, these models are converged on CpG islands of selected genes including ELOVL2 known to function in lipid synthesis relevant to cardiovascular health [4, 5, 11–13]. Altogether, these data highlight the fact that certain genomic regions may harbor DNA methylation with changes being mostly consistent with the progression of age, and that additional age-related CpG sites within the regions might be used to improve the accuracy of age prediction.

In the present study, we devised a discovery cohort by deliberately recruiting more aged human subjects than other age categories, aiming to increase the prediction accuracy by diminishing the effect of epigenetic drift. By employing an independent validation cohort and a multi-center test using trace blood samples taken directly from the forensic casework, we demonstrated that DNA methylation from a single genomic region is sufficient for the prediction of chronological age. We propose that our age-prediction model features substantially improved precision, ease of manipulation, bulk processing, and low cost.

## Material and methods

### The study population

This study was conducted in the Chinese Han ethnic group. We collected 319 blood samples aged 30-70 years from the community of Baoshan District, Shanghai, China, including 181 females and 138 males, with 46.4% aged over 60 years. All blood samples were collected in Blood Nucleic Acids Tubes (Thermo Fisher, catalog: 4342792) and stored at -80°C until use, avoiding repeated freezing and thawing of plasma to prevent DNA degradation and contamination. In addition, dried bloodstains from forensic casework were provided by the public security from Beijing (n=5), Yangzhou (n=59), and Shanghai (n=8). Written informed consent was obtained prior to sample collection from every participant after explaining the objectives and procedures of the study.

### DNA extraction

For isolation of total DNA, 300 μl of whole blood was mixed with 3 μl RNase A (200 ng/μl, ABclonal, Catalog: RM29870) and 20 μl Proteinase K Solution (Magen, catalog: D6310-03B) and incubated for 15 min at 37°C with shaking. Bloodstains were cut and mixed with 20 μl Proteinase K Solution and 400 μl Digestive Solution ATL (Magen, catalog: D6310-03B) and then processed according to the manufacturer’s instructions (Magen, catalog: D6310-03B). DNA concentration was measured using NanoDrop (Thermo Fisher).

### Methylation analysis

Unmethylated cytosine was converted to uracil using the Bisulfite Conversion Kit (Singlera, catalog: EP110192). 200 ng-1 μg of DNA was added with ddH_2_O to make up to 60 μl and then processed according to the manufacturer’s instructions (Singlera, catalog: EP110192). Primers were designed by the website https://amplicondesign.dkfz.de/, with degenerate sequence as followings (R=A/G;Y=C/T): ELOVL2-F:5’-TACACGACGCTCTTCCGATCTYGGTYGGGYGGYGATTTGTA-3’; ELOVL2-R: 5’-GACGTGTGCTCTTCCGATCTACCCACCRAAACCCAACTAT-3’.

The concentration of each primer was 100 μM. 4 μl each of ELOVL2-F/R were mixed with ddH_2_O to make up to 100 μl of primer mix. Library pre-construction was performed using KAPA2G Fast Multiplex Mix (Roche, catalog: 2GFMPXKB). PCR reactions were carried out in a total volume of 25 μl, containing10 ng of template DNA, 1μl of primer mix and 12.5μl of 2X KAPA2G Fast Multiplex Mix and ddH_2_O. The PCR program operated with an initial denaturation step of 5 min at 95□, amplification for 25 cycles (denaturation for 15 s at 95□, annealing for 15 s at 58□ and extension for 30 s at 72□), and a final extension for 5 min at 72□. The amplified pre-hybridized libraries were then purified using VAHTS DNA Clean Beads (Vazyme, catalog: N411-02) with a volume ratio of 0.9 for the first round of sorting (DNA Clean Beads: DNA) and 0.3 for the second round of sorting. The purified products were subjected to secondary amplification using TaKaRa Ex Taq (TaKaRa, catalog: RR53A) and adaptor-specific primer (F: AATGATACGGCGACCACCGAGATCTACACAGCGCTAGACACTCTTTCCCTACACGA CGCTCTTCCGATCT; R: CAAGCAGAAGACGGCATACGAGATAACCGCGGGTGACTGGAGTTCAGACGTGTG CTCTTCCGATCT). PCR reactions were carried out in a total volume of 25 μl, containing10 μl of purified products, 2.5μl of 10×Ex Taq Buffer, 2μl of dNTP, 1μl of TaKaRa Ex Taq, 1 µl each of adaptor-specific primer F/R and 8.4µl of ddH2O. The PCR program operated with an initial denaturation step of 5 min at 95□, amplification for 13 cycles (denaturation for 15 s at 95□, annealing for 15 s at 58□ and extension for 30 s at 72□), and a final extension for 5 min at 72□. VAHTS DNA Clean Beads was used for purification with a volume ratio of 0.7 for the first round of sorting (DNA Clean Beads: DNA) and 0.3 for the second round of sorting. The concentration of the final library was determined using Qubit 2.0 (Invitrogen) Libraries were sequenced on the Illumina NovaSeq 6000 system (paired end; 150 bp).

### High-throughput sequencing data analysis

All sequencing reads were processed with Trim Galore (v0.6.6) [14] with the parameters “--nextseq 30 --paired” to remove the adapter sequences (AGATCGGAAGAGC) from NovaSeq-platforms and reads longer than 20 bp were kept. Reads that passed the quality control procedure were kept and mapped to the Homo sapiens genome (GRCh38) using bismark (v0.24.1) [15] with default parameters. Uniquely mapped read pairs were extracted using samtools (v1.17) [16] Methylation level was extracted by bismark.

### Age prediction model

To develop an age prediction model, we employed elastic net regression. Age prediction was trained by regressing chronological age on methylation level using the discovery cohort (N = 191). To begin, we randomly split the discovery cohort into training (70%) and test (30%) sets with balanced ages. Model optimization including hyperparameter tuning was done by a grid search with leave-one-out cross-validation (LOOCV) based on training sets. Model performance was assessed on the test set, using several statistics including median absolute error (MAE), Pearson’s correlation coefficient and its associated *p* value. Furthermore, we performed a cross-validation scheme for arriving the least biased estimates of the accuracy of the aging clocks, consisting of leaving out a single sample from the regression, predicting age for that sample, and iterating over all samples on the discovery cohort. The best-tuned hyperparameter α was 0.01, and λ was 1.2. Above model training and hyperparameter tuning were performed with R packages caret (v6.0-93) and glmnet (v4.1-4).

## Results

### The rational design of discovery aging cohort and multiplexing assay

To establish a discovery cohort, we collected 191 peripheral blood samples from volunteers of Chinese Han ethnicity at the Baoshan District community of Shanghai. We deliberately recruited more aged subjects equal to or older than 65 years compared to other age categories (**Fig. 1A and Fig. S1A**). Our rationale lies at the fact that elderly individuals, due to medical history and age, may couple with epigenetic drifting that confounds DNA methylation status, such that only CpG sites that are mostly correlated with age could be selected. Different from previous reports that relied on low-throughput assay, we assessed DNA methylation from a single genomic region of ELOV2 gene by massive parallel DNA sequencing (**Fig. 1A**). We consolidated the feasibility of the analytical pipeline. Sequence analysis demonstrated sufficient read counts, with minimally more than 1,000 for the region involved (**Fig. S1B)**.

**Fig. 1.**
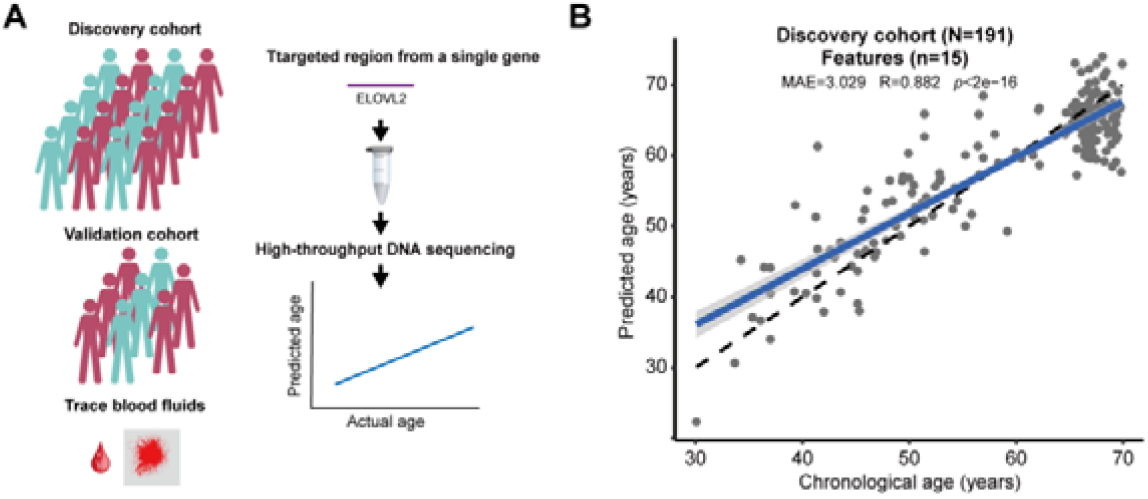
A new age-prediction model based on DNA methylation. (A) We constructed an age-prediction model with a discovery cohort, which was consolidated by an independent validation cohort and a single-blind test using trace blood samples relevant to forensic casework (left panel). (B) A new age-prediction model based on 15 CpG features was established using a discovery cohort consisting of 191 human subjects (MAE=3.209, R=0.882, *p*<2e-16).

### A new age-prediction model

We constructed a new age-prediction model using the discovery cohort and a pipeline based on DNA methylation from a single genomic region of ELOV2 and massive parallel DNA sequencing (**Fig. 1A**). We applied elastic net regression and cross-validation to construct a new age-prediction model that involved 15 CpG sites from ELOVL2 gene, with 13 new CpG features not being used by previous models (**Table 1**). This model demonstrated significantly improved accuracy with MAE as low as 3.029 (R=0.882, *p*<2e-16) (**Fig. 1B**).

**Table 1.**
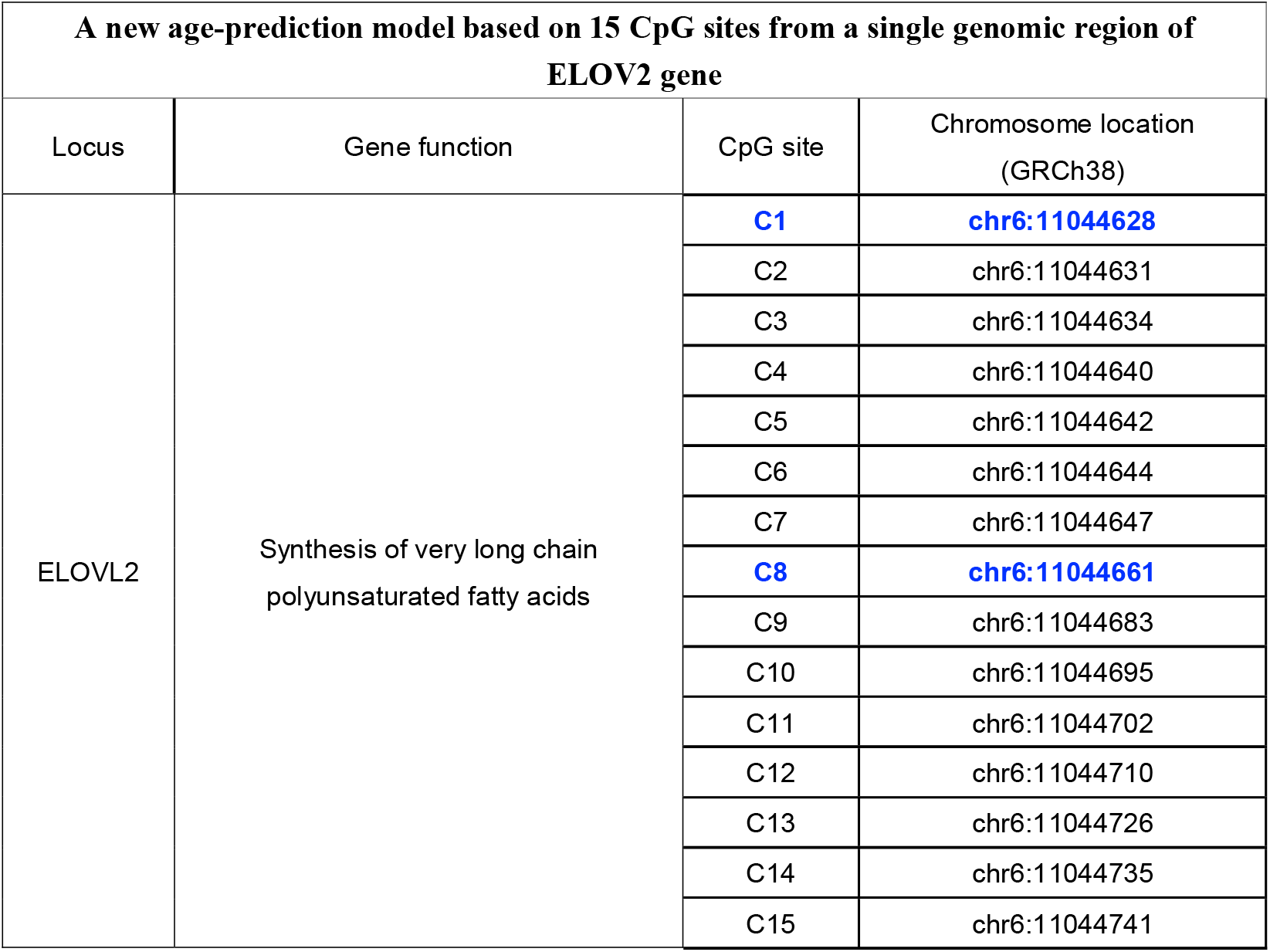
A new age-prediction model based on 15 CpG sites. A list of 15 CpG sites used in the age-prediction model. CpG sites used by previous models were highlighted in bold blue.

| A new age-prediction model based on 15 CpG sites from a single genomic region of ELOV2 gene |  |  |  |
| --- | --- | --- | --- |
| Locus | Gene function | CpG site | Chromosome location (GRCh38) |
| ELOVL2 | Synthesis of very long chain polyunsaturated fatty acids | **C1** | **chr6:11044628** |
|  |  | C2 | chr6:11044631 |
|  |  | C3 | chr6:11044634 |
|  |  | C4 | chr6:11044640 |
|  |  | C5 | chr6:11044642 |
|  |  | C6 | chr6:11044644 |
|  |  | C7 | chr6:11044647 |
|  |  | **C8** | **chr6:11044661** |
|  |  | C9 | chr6:11044683 |
|  |  | C10 | chr6:11044695 |
|  |  | C11 | chr6:11044702 |
|  |  | C12 | chr6:11044710 |
|  |  | C13 | chr6:11044726 |
|  |  | C14 | chr6:11044735 |
|  |  | C15 | chr6:11044741 |

### Validation by an independent cohort

We validated the new model in an independent cohort comprising 128 subjects. This experiment showed that our model could reach 2.663 for MAE (R=0.946, *p*<2e-16) (**Fig. 2A**). When the maximum difference between predicted and actual age was set as 5 years, we observed 73.4% successful predictions for the entire cohort (**Fig. 2B**), and notably 79.2% success for the age category between 50 and 60 (**Fig. 2B**).

**Fig. 2.**
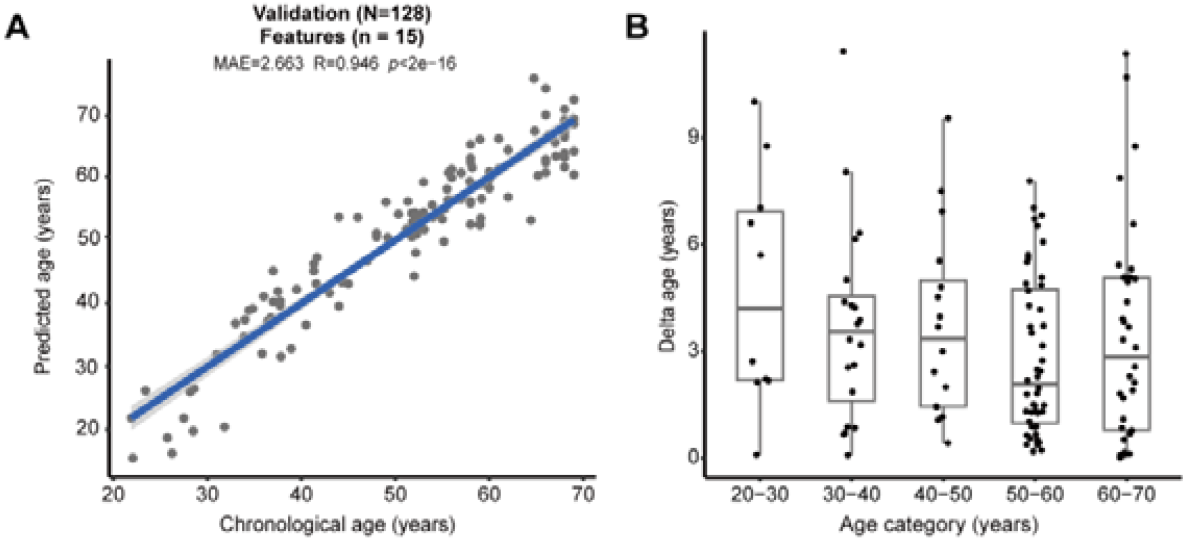
Validation by an independent cohort. (A) The validation cohort consisting of 128 human subjects confirmed the accuracy of the model (MAE=2.663, R=0.946, *p*<2e-16). (B) The accuracy of prediction in each age category of the validation cohort. Successful prediction could reach 73.4%, when the deviation between predicted versus actual age was set as 5 years (dashed lines).

### Efficacy to predict age from trace bloodstains

We attested the model with trace blood samples that were practically relevant to the crime scenes in the real world. We initiated a multi-center test using dried bloodstains from forensic casework provided by public security agencies from Beijing (n=5), Yangzhou (n=59), and Shanghai (n=8) (**Fig. 3A**). Without prior knowledge of age, our model could reach 3.039 for MAE (R=0.926, *p*<2.2e-16) (**Fig. 3A**). Notably, when the maximum difference between the predicted and actual age was set as 5 years, our data demonstrated 83.3% success (**Fig. 3B**). Significantly, this result illustrates the ability of our model of using DNA methylation sites from a single genomic locus, together with the streamlined analytical pipeline, to handle trace blood samples, strongly supporting its readiness in forensic application.

**Fig. 3.**
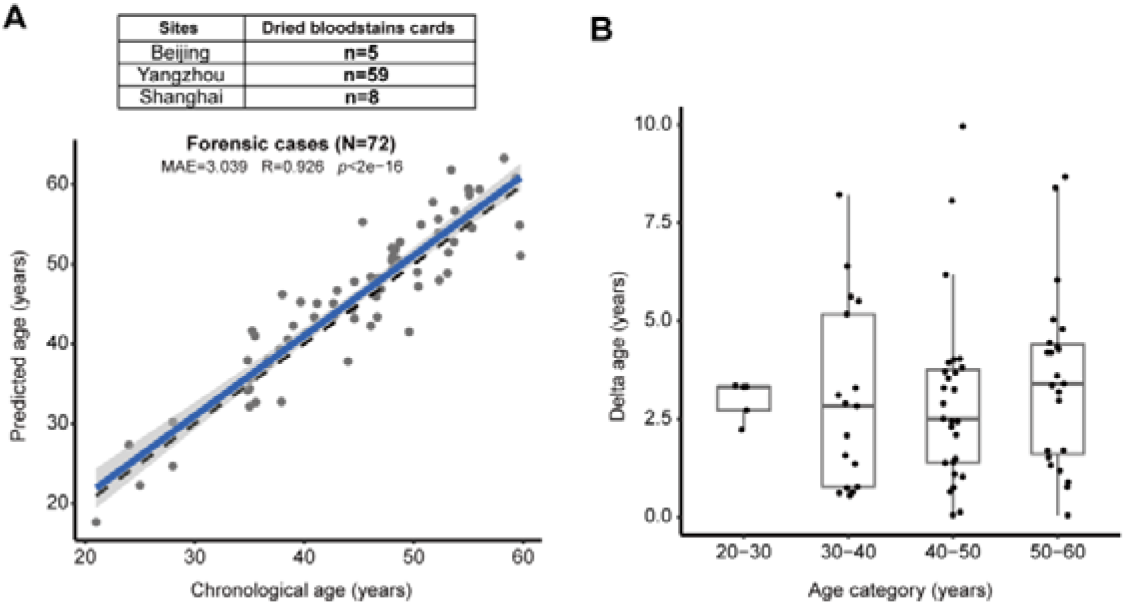
Validation by trace blood samples from forensic casework. (A) The validation cohort consisting of 72 dried bloodstains subjects confirmed the accuracy of the model (MAE=3.039, R=0.926, *p*<2e-16). (B) Successful prediction could reach 83.3% from bloodstains and trace blood fluids, when the deviation between predicted versus actual age was set as 5 years.

## Discussion

Estimation of age is informative in many forensic contexts, e.g., classically used in investigative leads for crime scenes [17]. The demand of this application is now rapidly growing in age categorization of illegal immigrants as well as asylum seekers in which context valid identification documents are usually missing [18]. Thus far, forensically practical age-prediction models have been established by the assessment of DNA methylation status, known as epigenetic clock [19–21]. The currently available models involve the use of 5-8 CpG sites and low-throughput analytical platforms, e.g., pyrosequencing [2, 3, 22, 23]. In the present study, we examine DNA methylation from a single genomic region of ELOV2 followed by high-throughput DNA sequencing. Trained by a carefully designed aging cohort, we propose a new epigenetic clock model that includes 15 age-related CpG sites.

Epigenetic DNA methylation is one of the hallmarks of aging, laying critical foundation for its application to estimate the chronological age of human subjects. However, it is well-known that stochastic changes in DNA methylation occur with age, smoking habit, alcoholic consumption, and disease situations [24–28]. Consequently, an age-related increase in interindividual variability and reciprocally a decline in accuracy have been reported in many age-prediction models based on biomarkers of DNA methylation [2, 3, 5, 29]. Hence, we designed a training cohort by deliberately recruiting more elderly subjects than other age categories, such that our model could be built with CpG sites that are mostly consistent with the progression of age. Presumably, the accuracy of age prediction could be substantially improved by increasing the number of DNA methylation loci. As such, we focus on ELOV2 gene therein repeatedly used by various early models [4, 5, 12, 13, 30, 31]. By high-throughput DNA sequencing, we can obtain profiles of all DNA methylation loci from which we construct a new model comprising 15 CpG sites. Despite increased number of DNA methylation loci, our pipeline has been significantly simplified, since only a single genomic region is employed for analysis. Moreover, the nature of our analytical pipeline, by measuring the ratios between methylated and non-methylated CpG from high-throughput sequencing, with minimally 1,000 read counts for each region, supports the consistency and robustness of the result. Empowered by this new model, we could predict age with a success rate of 73.4% (±5 years) in an independent validation cohort and a success rate of 83.3% (±5 years) in a multi-center test using dried bloodstains taken from real world forensic casework.

Compared to the early models based on low-throughput assay, we apply high-throughput sequencing, thus allowing bulk processing of large volume of samples. This ability is of paramount urgency, given the exponentially increasing casework of illegal immigrants and asylum seekers resulted from geographic conflict. Moreover, another issue to consider is cost. It is important to note that the cost for massive parallel DNA sequencing has been significantly reduced, especially feasible for batch testing. Taken together, we propose a new age-prediction model featuring substantially improved accuracy, ease of manipulation, ability for bulk processing, and low cost.

## Conclusion

In this study, we propose a new age-prediction model by assessment of DNA methylation from a single genomic region of ELOV2, which has substantially improved accuracy, ease of manipulation, ability for bulk processing, and low cost. To conclude, this model can be readily applied in both classic and newly emergent forensic contexts that require the estimation of chronological age.

## Acknowledgments

This work was supported by grants from the National Natural Science Foundation of China (8237020490) to L.Q., innovative clinical research project of the Chang Zheng Hospital, Naval Medical University (2020YLCYJ-Z09) to L.Q., clinical research projects initiated by investigators in exemplary research award (2023YJBF-FH04) to L.Q., the Shanghai Key Laboratory of Aging Studies (19DZ2260400) to N.L., the National Key Research and Development Project of China (2018YFC2000203, 2018YFC2000204) and to M.S., and an open project grant of Institute of Forensic Science of China (2021FGKFKT02) to X.Y..

## Disclosure of potential conflicts of interest

The authors declare no competing interests.

## Research involving human participants and/or animals

All procedures performed in studies involving human blood samples were in concordance with the ethical standards of the institutional research committee and with the 1964 Helsinki Declaration and its later amendments or comparable ethical standards. This article does not contain any studies with animals performed by any of the authors.

## Ethics approval

The project was approved by the Medical Ethics Committee of Shanghai Chang Zheng Hospital (2022SL006) and the Third Research Institute of Ministry of Public Security of China (2024SH003). Informed consent was obtained from all subjects in accordance with the local research Ethics Committee guidelines.

## Informed consent

Written informed consent was obtained prior to sample collection from every participant after explaining the objectives and procedures of the study.

## Data Availability Statements

All high-throughput DNA methylation data have been deposited at the Gene Expression Omnibus under accession number GSE253549 (secure token for reviewers: ezgdoyaqrlexnyx).

## Notes

### Competing Interest Statement

The authors have declared no competing interest.

