## Supplementary figures and images for "Assessment of DNA methylation from a single genomic region of ELOV2 is sufficient to predict chronological age"

### Supplemental Figure 1

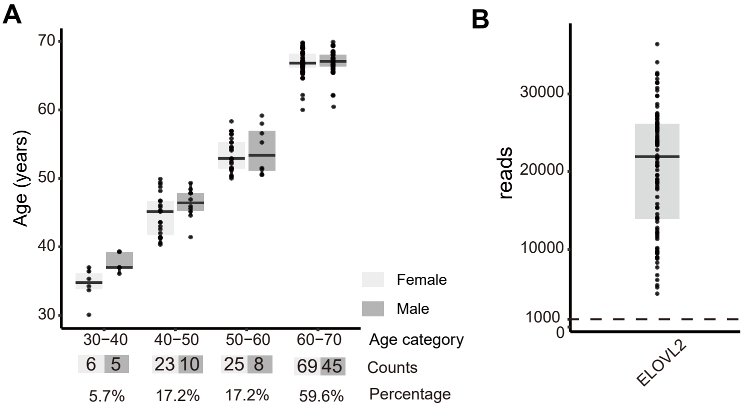
